# Sex and age don’t matter but breed type does - Factors influencing eye wrinkle expression in horses

**DOI:** 10.1101/567149

**Authors:** Lisa Schanz, Konstanze Krueger, Sara Hintze

## Abstract

Identifying valid indicators to assess animals’ emotional states is a critical objective of animal welfare science. In horses, eye wrinkles caused by the contraction of the inner eyebrow raiser have been shown to be affected by pain and other emotional states. Whether individual characteristics of a horse systematically affect eye wrinkle expression has not yet been studied. Therefore, the aim of this study was to assess how age, sex, breed type, body condition and coat colour affect the expression and/or the assessment of eye wrinkles in horses. To this end, we adapted the eye wrinkle assessment scale from Hintze et al. (2016) and assessed eye wrinkle expression on pictures taken from the left and the right eye of 181 horses in a presumably neutral situation, using five outcome measures: a qualitative first impression reflecting how worried the horse looks, the extent to which the eyebrow is raised, the number of wrinkles, their markedness and the angle between a line through the eyeball and the topmost wrinkle. All measures could be assessed highly reliable with respect to intra- and inter-observer agreement. Breed type affected the width of the angle (F_2, 114_ = 8.20, p < 0.001), with thoroughbreds having the narrowest angle (*M* = 23.80, *SD* = 1.60), followed by warmbloods (*M* = 28.00, *SD* = 0.60), and coldbloods (*M* = 31.00, *SD* = 0.90). None of the other factors affected any of the outcome measures, and eye wrinkle expression did not differ between the left and the right eye area (all p-values > 0.05). Consequently, horses’ characteristics age, sex and coat colour did not systematically affect eye wrinkle expression, whereas ‘breed type’ explained some variation in ‘angle’; how much eye wrinkle expression is affected by emotion or perhaps mood needs further investigation and validation.

## 1 Introduction

Assessing emotional states in animals is a critical goal in animal welfare science, but it is generally agreed that the subjective experience of an emotion cannot be assessed directly (but see Wemelsfelder, 1997 for a different point of view). However, emotional states are multifaceted, including not only the subjective experiences but also behavioural, physiological and cognitive components, which can be assessed objectively and could therefore serve as indicators to infer animals’ subjective experiences (e.g. Paul et al., 2005). Ideally, such indicators can be assessed non-invasively as well as reliably across various contexts, and do not require the animals to be trained. Spontaneous behaviour and behavioural expressions, including facial expressions, are promising examples of indicators to assess animals’ emotional states.

Facial expressions in animals have mainly been studied using Facial Action Coding Systems (FACSs) and Grimace Scales (GSs). In FACSs, all possible facial muscle movements and resulting expressions are systematically catalogued as Action Units or Action Descriptors (Ekman and Friesen, 1976). Originally developed for humans, FACSs have now been adapted to different animal species, including primates (orang-utans: Caeiro et al., 2013; macaques: Parr et al., 2010, chimpanzees: Parr et al., 2007a, 2007b; gibbons: Waller et al., 2012), dogs (Waller et al., 2013), cats (Caeiro et al., 2017) and horses (Wathan et al., 2015). Besides their application in comparative psychology (e.g. Vick et al., 2007) and research on the evolution of emotional communication (e.g. Parr and Waller, 2006), FACSs have more recently also been used to associate facial expressions with emotional states (Caeiro et al., 2017).

Grimace Scales have been developed for the specific assessment of pain by comparing the facial expressions of animals in painful and painless conditions and systematically identifying the resulting changes. They exist for a range of species, including laboratory animals (e.g. mice: Langford et al., 2010; rats: Sotocinal et al., 2011; rabbits: Keating et al., 2012), farm animals (e.g. dairy cows: Gleerup et al., 2015; sheep: McLennan et al., 2016; including lambs: Guesgen et al., 2016; piglets: Di Giminiani et al., 2016; and horses: Dalla Costa et al., 2014). In horses, two scales, namely the Horse Grimace Scale (HGS, Dalla Costa et al., 2014) and the Equine Pain Face (EPF, Gleerup et al., 2015) have been published. These scales have been developed by comparing the facial expressions of horses before and after castration while undergoing different pain treatments (HGS) and horses exposed to two noxious stimuli expected to induce pain (EPF). Both studies identified, beyond several other Action Units, eye wrinkles above the horses’ eyes as more prevalent when horses were in pain (HGS: ‘tension above the eye area’, EPF: ‘angled eye’). In addition to their association with pain, eye wrinkles are often dubbed ‘worry wrinkles’ in the equine community, suggesting a more general association with negative emotions. To investigate whether eye wrinkle expression is affected by emotional states, Hintze and colleagues (2016) developed a detailed scale of this expression caused by contraction of the inner eyebrow raiser (levator anguli oculi medialis and corrugator supercilii muscles, identified as Action Unit 101 in EquiFACS; Wathan et al., 2015). This scale describes different characteristics of the wrinkles, including the number and markedness of wrinkles and the angle between a horizontal line through the eyeball and the topmost wrinkle. The scale was applied to pictures of sixteen horses, which were exposed to two presumably positive situations (grooming, food anticipation) and two presumably negative situations (waving of plastic bag, food competition) in a counterbalanced order. It was found that the angle decreased during grooming (muscle relaxation) and increased during food competition (muscle contraction) compared to control phases but no other characteristics of the eye wrinkle expression were systematically affected by these situations.

The results of this first study on the association between eye wrinkle expression and horses’ emotional states are promising with respect to the angle measure but further validation is needed before eye wrinkles can be used as a potential indicator of emotional valence in horses. Moreover, it needs to be investigated whether individual characteristics of horses, e.g. their age, sex, breed type or body condition, affect eye wrinkle expression systematically. Some of these factors have been shown to influence facial wrinkles, including eye wrinkles, in other species. Facial wrinkles, which are not caused by muscle movement, have, for example, been used to identify individual white rhinos (Patton and Campbell, 2011) and to determine age in humans (Akazaki et al., 2002; Kwon and da Vitoria Lobo, 1999). The wrinkles assessed in these studies may have a genetic basis or may have been affected by other aspects, for example reduced skin tension with increased age in humans (Akazaki et al., 2002). Even though eye wrinkles in horses differ from the described facial wrinkles as they are caused by muscle contraction, they might still be systematically affected by individual characteristics. Moreover, factors that may not affect the expression itself, but the assessment of the expression need to be considered. Coat colour may be such a factor affecting the visibility of single wrinkles. According to Dalla Costa et al. (2014), for example, it is easier to score lighter horses compared to darker horses for Action Units of the Horse Grimace Scale. Additionally, it is important to know – for the interpretation of the assessment - whether eye wrinkle expression differs between the left and the right eye area of a horse. So far, there has been limited research on facial asymmetry in horses. One study showed that horses that were groomed in a gentle manner (assumed to elicit positive emotions) showed asymmetric ears less often than horses groomed in a standard procedure (assumed to elicit negative emotions; Lansade et al., 2018). If any of the aforementioned characteristics systematically affects eye wrinkle expression or its assessment, it needs to be considered when using eye wrinkles as a potential indicator of horses’ emotional states. Consequently, we investigated how age, sex, breed type, body condition and coat colour affect the expression and/or assessment of eye wrinkles in horses, and whether the expression differs between the area above the left and right eye. To this end, we adapted the eye wrinkle assessment scale from Hintze et al. (2016) to assess the individual characteristics on eye wrinkle expression in pictures of the left and right eye of 181 horses in a ‘neutral’ situation.

## 2 Animals, material and methods

### 2.1 Animals and Housing

This study included 181 horses (70 mares, 62 geldings and 49 stallions) of varying age and breed (SI Table 1). The horses’ age ranged from 4 months to 28 years (mean (*M*) = 11.94, standard deviation (*SD*) = 6.48) and 44 different breeds and crossbreeds were included. Each horse was assigned a ‘breed type’ (coldblood, warmblood, thoroughbred, pony) according to the breeds’ stud books, resulting in 36 coldbloods, 104 warmbloods, 17 thoroughbreds, 7 ponies and one horse with unknown breed.

Horses were derived from seven farms across three countries: five farms in Germany (Farms 1 – 5), one farm in the United States of America (Farm 6), and one farm in Switzerland (Farm 7). Housing conditions varied between and within farms with horses kept either in standard single boxes, single paddock boxes or in groups. Horses in standard single boxes were kept on wood shavings, straw or a mixture of both with visual and in some cases physical contact to conspecifics. Horses living in a paddock box were also kept individually but with more space (box plus paddock) and physical contact to conspecifics was always possible. All horses kept in boxes were turned out either on paddocks or pastures, depending on the weather, in groups of at least two, except for stallions on Farm 7 and two older stallions on Farm 6, which were turned out individually. When kept in groups, horses were housed either in a pen system or on pasture, each with a shelter and a bedded lying area. Horses on all farms were fed hay and concentrates with the number of feedings per day varying between farms (hay: ranging from one feeding a day to hay *ad libitum*, concentrates: ranging from one to three feedings). On all farms, *ad libitum* access to water was provided by automatic drinkers, except for Farm 6, where water was supplied in buckets in the stables and big tanks on pastures. All horses were either exercised (riding, carriage driving), turned out on paddock or pasture, longed, or walked in a horse walker daily.

### 2.2 Data collection

#### 2.2.1 Pictures taken from the eye area

Data were collected on Farms 1 – 6 during spring and summer 2015, and on Farm 7 in summer 2014 and spring 2016. Horses were photographed in a presumably neutral situation in their habitual environment between 9 AM and 5 PM. If any disturbance (visual or acoustic) occurred, photographing was stopped until the disturbance subsided. The photographer stood at a 45° angle to each horse’s head, while an additional person was loosely holding the horse’s halter. Both were instructed to interact as little as possible with the horse to keep the influence of handling to a minimum. Several pictures of the horses’ left and right eye areas were taken with a Canon G1X camera by one photographer (Farms 1 - 6) and with a Nikon D200 with professional lenses (Nikon AF Micro Nikkon 60mm f/2.8D and Nikon 80-200mm f/2.8 AF-D telephoto lens) by two photographers (Farm 7).

#### 2.2.2 Body Condition Score (BCS)

After pictures from both eyes were taken, the body condition of each horse was assessed by visual and tactile evaluation using the scale developed by Henneke et al. (1983). With this scale, the presence or absence of adipose tissue and the visibility of bone structures is assessed on a nine-point scale (1 – 9, half points can be given). One person assessed the body condition of horses on all farms except for Farm 7, where the assessment was done by another person with the same scale. Sixteen horses on Farm 7 were not assessed. The BCS ranged from 2.5 to 8 (*M* = 5.3, *SD* = 0.9).

### 2.3 Picture processing

All pictures were screened and blurry pictures and pictures on which the eye area was not fully visible were excluded from further assessment. Pictures were defined as blurry if wrinkles were not clearly detectable or the beginning and/or end of wrinkles was not visible (Hintze et al., 2016). From the remaining pictures, two or three pictures per horse and eye (left, right) were randomly selected for scoring (62 horses x 4 pictures and one horse x 2 pictures, data collection in summer 2014; 118 horses x 6 pictures, data collection in spring/summer 2015 and spring 2016) using the ‘sample’ function in R (R Version 3.5.1, R Studio Version 1.1.453) and resulting in a total of 958 pictures. The selected pictures were cropped to only show the eye area needed for scoring and picture size was standardised using Microsoft Picture Manager (version 2018.18051.17710.0).

### 2.4 Eye wrinkle assessment scale

In the present study, we used an adapted version of the eye wrinkle assessment scale developed by Hintze et al. (2016) resulting in five outcome measures: ‘qualitative assessment’, ‘eyebrow raised’, ‘number’, ‘markedness’ and ‘angle’ (for definitions and scoring details see Fig. 1). ‘Number’ (C), ‘markedness’ (D) and ‘angle’ (E) were defined as previously suggested by Hintze and colleagues (for a direct comparison of the two scales see Table 1). For the outcome measure ‘qualitative assessment’ (A), we adapted the definition by Hintze et al. (2016): The original definition focused on three particular aspects of the eye wrinkle expression, namely “the number […], markedness, and the angle” (Hintze et al., 2016, p. 6), whereas in the present study we aimed to better capture the overall first subjective impression with respect to how ‘worried’ the horse actually looks (‘not worried’ to ‘extremely worried’) without focusing on other aspects at this stage. Moreover, instead of using an ordinal scale with three distinct categories for ‘qualitative assessment’, we used Visual Analogue Scales (VAS) to possibly get more sensitive measures (Tuyttens et al., 2009). VAS are an instrument to measure variables that range across a continuum of values (Wewers and Lowe, 1990). It is presented as a horizontal line with the end anchors labelled as the boundaries of this variable (e.g. ‘not worried’ to ‘extremely worries’). For each variable (e.g. ‘worriedness’) a vertical mark through the line is placed at a position, which is deemed appropriate by the observer (Wewers and Lowe, 1990). We used the freely available programme AVAS (Adaptive Visual Analog Scales), which stores the positions of the mark on the line in an excel sheet (Marsh-Richard et al., 2009). VAS were also used to assess the outcome measure ‘eyebrow raised’ (B), which replaced ‘eyelid shape’ used in the scale described by Hintze et al. (2016). Inter-observer agreement for ‘eyelid shape’ was only moderate in the study by Hintze et al. (2016). During discussions aiming to improve the definition and thereby inter-observer agreement, we realised that we could capture the amount the skin above the eye (brow region in humans) is pulled in the medio-nuchal direction (resembling Action Unit 1 in human FACS, Wathan et al., 2015) more easily than the shape of the eyelid and consequently replaced ‘eyelid shape’ with what we named ‘eyebrow raised’ (‘not raised’ to ‘strongly raised’). Moreover, we excluded the binary outcome measures ‘eye white’ used by Hintze and colleagues (2016) to assess the presence or absence of visible sclera. Since our study focused on the effect of horses’ characteristics on eye wrinkle expression in neutral situations and not on general changes in the eye area caused by different emotional situations, we decided to drop this measure.

**Fig. 1.**
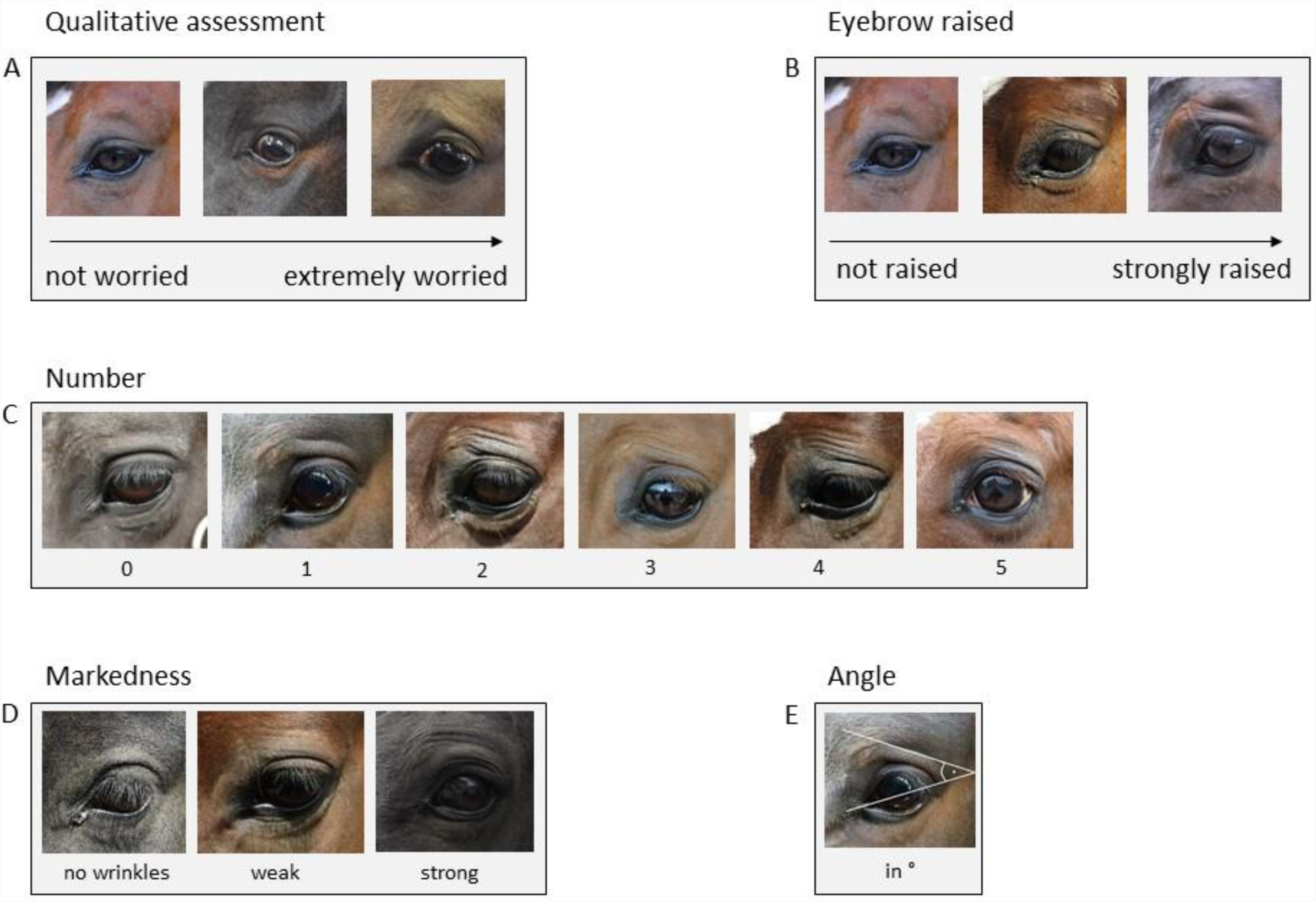
Eye wrinkle assessment scale. (adapted from Hintze et al. 2016). (A) Qualitative assessment: The overall first subjective impression of the eye area with respect to how ‘worried’ the horse actually looks as assessed on a Visual Analogue Scale ranging from ‘not worried’ to ‘extremely worried’. (B) Eyebrow raised: The amount the skin above the eye (brow region in humans) is raised, assessed on a Visual Analogue Scale ranging from ‘not raised’ to ‘strongly raised’. (C) Number: Only wrinkles above the eye and those of a minimum length of one third of the eyeball’s diameter are considered. A deep indent, often seen in older horses, is not considered as a wrinkle (as it is not caused by muscle contraction of the inner eyebrow raiser). Moreover, wrinkles originating on the eyelid are not counted. (D) Markedness: The depth and width of the wrinkles is assessed. If the markedness differs between wrinkles, the most prominent wrinkle is assessed. ‘No wrinkle’: no wrinkle visible. ‘Weak’: wrinkles are flat and narrow lines. ‘Strong’: wrinkles are pronounced in depth and width. (E) Angle: The degree of the angle is measured on the intersection of the extension of a line drawn through the eyeball and the extension of the topmost wrinkle. The line through the eyeball extends from the medial to the lateral corner of the eyeball. If the medial corner is not clearly defined, the line goes through the middle of the tear duct.

**Table 1.**
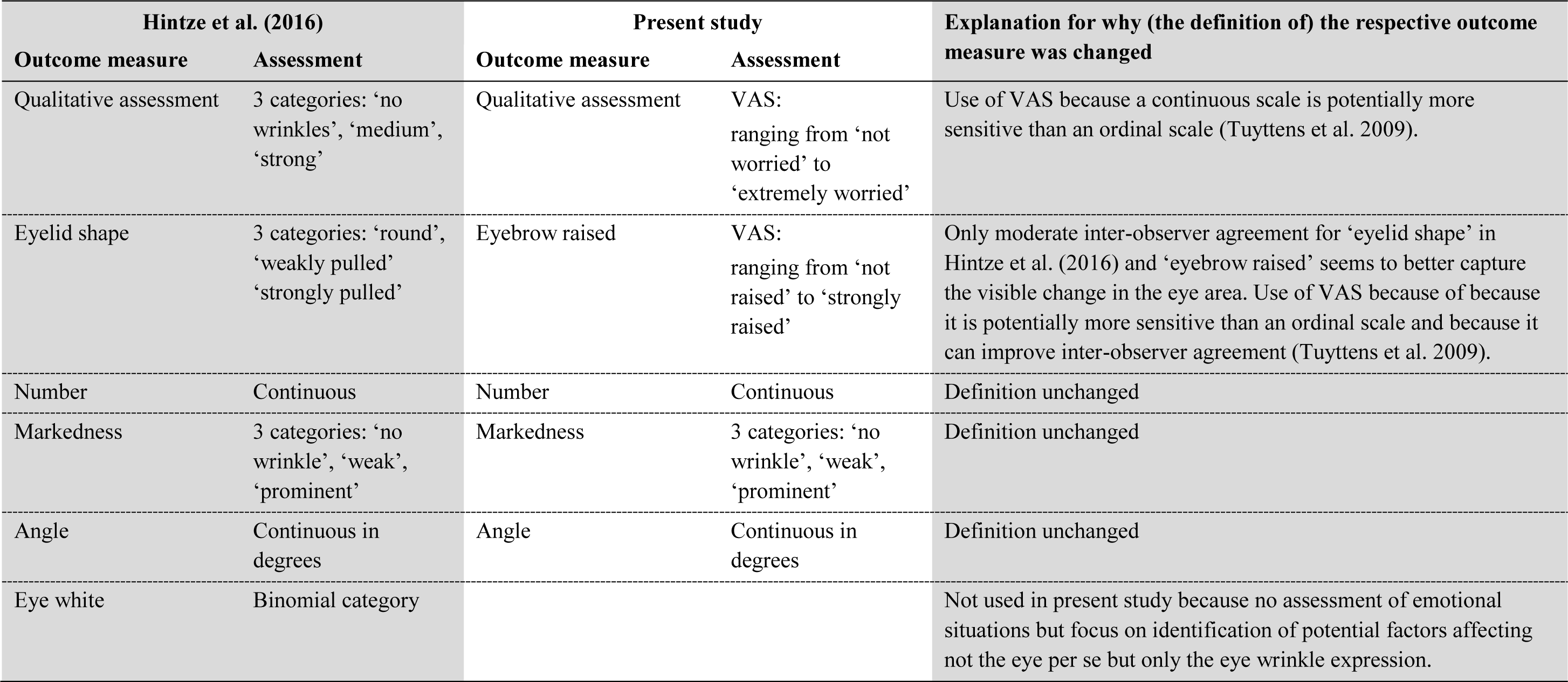
Comparison between the outcome measures used in the study by Hintze et al. (2016) and in the present study.

| Hintze et al. (2016) |  | Present study |  | Explanation for why (the definition of) the respective outcome measure was changed |
| --- | --- | --- | --- | --- |
| Outcome measure | Assessment | Outcome measure | Assessment |  |
| Qualitative assessment | 3 categories: ‘no wrinkles’, ‘medium’, ‘strong’ | Qualitative assessment | VAS: ranging from ‘not worried’ to ‘extremely worried’ | Use of VAS because a continuous scale is potentially more sensitive than an ordinal scale (Tuytens et al. 2009). |
| Eyelid shape | 3 categories: ‘round’, ‘weakly pulled’ ‘strongly pulled’ | Eyebrow raised | VAS: ranging from ‘not raised’ to ‘strongly raised’ | Only moderate inter-observer agreement for ‘eyelid shape’ in Hintze et al. (2016) and ‘eyebrow raised’ seems to better capture the visible change in the eye area. Use of VAS because of because it is potentially more sensitive than an ordinal scale and because it can improve inter-observer agreement (Tuytens et al. 2009). |
| Number | Continuous | Number | Continuous | Definition unchanged |
| Markedness | 3 categories: ‘no wrinkle’, ‘weak’, ‘prominent’ | Markedness | 3 categories: ‘no wrinkle’, ‘weak’, ‘prominent’ | Definition unchanged |
| Angle | Continuous in degrees | Angle | Continuous in degrees | Definition unchanged |
| Eye white | Binomial category |  |  | Not used in present study because no assessment of emotional situations but focus on identification of potential factors affecting not the eye per se but only the eye wrinkle expression. |

### 2.5 Scoring

All 958 pictures were scored in random order. We first scored the outcome measure ‘qualitative assessment’ (assessed on a VAS) on all pictures to capture the first subjective impression of the eye wrinkles without being biased by the other measures. We then continued with assessing ‘eyebrow raised’ (also assessed on a VAS), followed by ‘number’, ‘markedness’ and ‘angle’ (measured with CorelDraw, Corel Cooperation, 2014).

### 2.6 Intra- and inter-observer agreement

To assess both intra- and inter-observer agreement, a sample of all pictures was re-scored by the same rater (to assess intra-observer agreement) and by a second rater (to assess inter-observer agreement). To assess intra-observer agreement, ten out of each subset of 50 pictures (in total n = 192 pictures) were randomly selected and scored a second time, at the earliest the day after the first scoring. To assess inter-observer agreement, a sample of 10 % of all pictures (n = 96 pictures) was scored by rater B after rater A had finished scoring.

### 2.7 Coat colour

The horses’ ‘coat colour’ was assessed by inspecting the area above the eye relevant for the eye wrinkle assessment on each picture. One of the following colours was assigned to each picture: black/dark bay, medium bay, light bay, grey/white or undefinable (e.g. more than one colour present in the relevant area). Both raters assessed all pictures together.

### 2.8 Ethical considerations

The ethical guidelines of the International Society for Applied Ethology were respected while carrying out this experiment. For photographing, horses were loosely held on a halter (a normal routine for all horses used in this study) without any further manipulation. Horses from Farm 7 were additionally used in two larger studies, which were approved by the Cantonal Veterinary Office in Vaud, Switzerland (license numbers 2804 and 2804_1).

### 2.9 Statistical analysis

#### 2.9.1 Intra- and inter-observer agreement

Both intra- and inter-observer agreement were analysed using the statistical programming language R (R version 3.5.1, R Core Team, 2014; RStudio version 1.1.453, RStudio Team, 2016). For continuous outcome measures, we calculated the Intraclass Correlation Coefficient (ICC, function: icc, package: irr, Gamer et al., 2012) to assess agreement between first and second scoring of rater A (intra-observer agreement) as well as between rater A and B (inter-observer agreement). For both intra- and inter observer agreement, we used a two-way mixed model with ‘single rater’ as ‘type’ and assessing absolute agreement (Koo and Li, 2016). A p-value lower than 0.05 and an ICC-value above 0.75 were considered ‘excellent agreement’ (Cicchetti, 1994). Lower and upper bounds of the 95 % confidence interval (CI) were assessed as a measure of deviation of the ICC. For the categorical outcome measure ‘markedness’, we calculated the association between two scorings with Cohen’s Kappa (function: kappa2, package: irr, Gamer et al., 2012), considering a p-value lower than 0.05 and a *κ*-value above 0.8 as ‘almost perfect agreement’ (Landis and Koch, 1977).

#### 2.9.2 Collinearity between explanatory variables

In a statistical model, collinearity of explanatory variables can affect model interpretation and increase the standard errors of the coefficients (Tu et al., 2005). To explore the relationship between the different explanatory variables (‘age’, ‘sex’, ‘Body Condition Score’, ‘breed type’, ‘coat colour’), each combination of them was tested for independence. When testing two categorical variables (e.g. ‘sex’ and ‘breed type’), a Crammer’s V test (square root of the fraction of Chi-square and the normalising factor) was performed, with V ranging between 0 and 1 (the closer to 0, the weaker the association; Cramér, 2016). We considered values above 0.7 as a strong association, indicating that only one variable should be included for further analyses. When testing one categorical and one continuous variable (e.g. ‘sex’ and ‘age’), a Kruskal-Wallis test was run to test whether the values of the continuous variable differed with respect to the different levels of the categorical variable. The Kruskal-Wallis test was run (instead of a one-way ANOVA) because both of our continuous variables (age, BSC) were not normally distributed. If the result of the Kruskal-Wallis test was significant (p ≤ 0.05), a pairwise Wilcoxon rank sum test was run as a post-hoc test to identify the levels of the categorical variable that differed from each other. If the results of the Wilcoxon rank sum test were statistically significant at all levels (p ≤ 0.05), only one variable was chosen for further analyses. This was the case for the association between ‘BCS’ and ‘breed type’, as the mean ‘BCS’ differed significantly across ‘breed types’ (Kruskal-Wallis trest: χ^2^_2_ = 145.99, p < 0.001; pairwise Wilcoxon rank sum test: all three p-values < 0.01) with coldbloods having the highest BCS (*M* = 6.00, *SD* = 0.70), followed by warmbloods (*M* = 5.10, *SD* = 0.70), and thoroughbreds (*M* = 4.80, *SD* = 0.90) having the lowest BCS. Additionally, ‘BCS’ and ‘sex’ were associated (Kruskal-Wallis test: χ^2^_2_ = 95.18, p < 0.001; pairwise Wilcoxon rank sum test: all three p-values < 0.01) with stallions (*M* = 5.90, *SD* = 0.90) having a greater ‘Body Condition Score’ than both mares (*M* = 5.20, *SD* = 0.80) and geldings (*M* = 5.10, *SD* = 0.70). Since ‘BCS’ had been assessed by two experimenters without testing for inter-observer agreement, we kept ‘breed type’ and ‘sex’ as the more reliable variables and excluded ‘BCS’ from all further analyses.

#### 2.9.3 Association between outcome measures

All outcome measures (‘qualitative assessment’, ‘eyebrow raised’, ‘number’, ‘markedness’, ‘angle’) were tested for association. If two outcome measures are strongly associated, statistical models should only be run for one (while the results can to some extent be transferred). The Kruskal-Wallis test described above was run to test for associations between ‘markedness’ and all continuous measures. As with the explanatory variables, only one outcome measure was retained for further analyses if the post-hoc pairwise Wilcoxon rank sum test was statistically significant for all levels of the categorical measure ‘markedness’. This was the case for the associations of ‘markedness’ with ‘qualitative assessment’ (χ^2^_2_ = 301.92, p < 0.001), ‘eyebrow raised’ (χ^2^_2_ = 333.06, p < 0.001) and ‘number’ (χ^2^_2_ = 881.66, p < 0.001); p-values < 0.01 for all pairwise Wilcoxon rank sum tests. Since ‘markedness’ was a categorical measure on an ordinal scale, it was probably the least sensitive measure, and we therefore dropped it. All remaining outcome measures were continuous variables, and a Pearson’s product-moment correlation coefficient was computed. If the correlation coefficient was greater than 0.7, indicating a ‘strong correlation’ (Martin and Bateson, 2007), only one outcome measure was selected for further analyses. A strong positive correlation between the outcome measures ‘qualitative assessment’ and ‘eyebrow raised’ was found (r = 0.9, p < 0.001), and we selected ‘qualitative assessment’ for all further analyses. All models were run for ‘qualitative assessment’, ‘number’ and ‘angle’.

#### 2.9.4 The effect of the different explanatory variables on the outcome measures

Model selection and execution was done in R. We employed a stepwise backwards selection procedure (functions: train, trainControl, packages: leaps, Lumley, 2017; caret, Kuhn et al., 2018; MASS, Venables and Ripley, 2002) based on the Akaike Information Criterion (AIC) to identify potentially relevant explanatory variables (‘age’, ‘sex’, ‘breed type’ and ‘coat colour’). The stepwise backwards selection procedure indicates which variables are included in the best fitting model (André et al., 2003; Broadhurst et al., 1997). We decided to include all categorical variables in the final model from which at least one level was selected as potentially relevant. After selecting all potentially relevant explanatory variables, we analysed the effect of these variables on the continuous outcome measures (‘qualitative assessment’, ‘number’ and ‘angle’) by running linear mixed-effects models (function: lme, package: nlme, Pinheiro et al., 2018). Explanatory variables selected as potentially relevant by the stepwise backwards selection procedure were included as fixed effects in the respective models, while random effects were included in all models as ‘eye’ nested in ‘horse’ nested in ‘farm’ (see Table 2 for an overview of all effects). To verify model assumptions, the residuals for each executed model were visually checked for normal distribution and homogeneity of variance. No transformation of the data was necessary. Post-hoc tests were performed with the function lsmeans (package: lsmeans, v. Lenth, 2016) using the ‘tukey method’ to correct for multiple testing.

**Table 2.** Overview of all fixed and random effects, whether they were treated as continuous or as categorical variable, as well as their ranges (mean (*M*), standard deviation (*SD*)) for continuous variables and their levels for categorical variables.

| Effect | Fixed or random | Type of variable | Range (*M*, *SD*) / Levels |
| --- | --- | --- | --- |
| Age | fixed (if selected) | continuous | 4 months – 28 years (*M* = 11.90, *SD* = 6.50) |
| Sex | fixed (if selected) | categorical | 3 levels (mare, gelding, stallion) |
| Breed type | fixed (if selected) | categorical | 4 levels (coldblood, warmblood, thoroughbred, pony) |
| Coat colour | fixed (if selected) | categorical | 4 levels (black/dark bay, medium bay, light bay, grey/white) |
| Farm | random | categorical | 7 levels (Farms 1 – 7) |
| Horse | random | categorical | 181 levels (horses 1 – 181) |
| Eye | random | categorical | 2 levels (left, right) |

#### 2.9.5 Side effects

To test for a possible difference between the right and the left eye, data were averaged per eye and horse by calculating the mean score of the respective pictures (function: aggregate, base R, R Core Team, 2014). We then ran a linear mixed-effects model (function: lme, package: nlme, Pinheiro et al., 2018) for all outcome measures with ‘eye’ as fixed effect and ‘horse’ nested in ‘farm’ as random effects. Model assumptions were verified by visually checking residuals for normal distribution and homogeneity of variance. No transformation of the data was necessary.

## 3 Results

### 3.1 Sample size and data structure

In total 958 pictures were assessed, including 118 horses with six pictures (three per eye), 62 horses with four pictures (two per eye) and one horse with only two pictures. Some pictures in the final sample could not be assessed reliably for all outcome measures due to low quality, leading to missing values for ‘qualitative assessment’ (n = 2), ‘number’ (n = 6), and ‘angle’ (n = 6).

In the final sample, ‘qualitative assessment’ ranged from 0.25 to 100 mm on the VAS (*M* = 36.63, *SD* = 31.45), and the ‘number’ of wrinkles varied from 0 to 5 wrinkles (*M* = 0.80, *SD* = 1.10). The ‘angle’ ranged from 5.8° to 50.6° (*M* = 13.00, *SD* = 14.70).

For one outcome measure (‘angle’) and two explanatory variables (‘breed type’, ‘coat colour’), a subset of the full data set was used. For ‘angle’ only pictures with at least one wrinkle (‘number’ ≥ 1) were included since an angle could only be measured if at least one wrinkle was identified, resulting in 427 pictures. For the analyses of ‘breed type’ on the different outcome measures, all ponies (n = 7) and the horse without known breed were removed from the data set due to the small sample size, leading to a remaining sample of 916 pictures. For 902 pictures a ‘coat colour’ could be assigned and were therefore used for subsequent analyses.

### 3.2 Intra- und inter-observer agreement

In this section results are given for all outcome measures including ‘eyebrow raised’ and ‘markedness’ in case these outcome measures will be included in future studies. Comparison of first and second scoring of rater A (intra-observer agreement) exceeded 0.9 for all continuous outcome measures (‘qualitative assessment’: ICC _agreement_ = 0.90, with a 95 % CI from 0.87 to 0.93; ‘eyebrow raised’: ICC _agreement_ = 0.94 with a 95 % CI from 0.92 to 0.96; ‘number’: ICC _agreement_ = 0.97, with a 95 % CI from 0.96 to 0.98; ‘angle’: ICC _agreement_ = 0.97, with a 95 % CI from 0.96 to 0.98) and 0.8 for the categorical outcome measure (‘markedness’: κ = 0.92) with all p-values being highly significant (p < 0.001). Inter-observer agreement was slightly lower than within rater A, but still exceeded 0.75 for all continuous outcome measures (‘qualitative assessment’: ICC _agreement_ = 0.80, with a 95 % CI from 0.70 to 0.87; ‘eyebrow raised’: ICC _agreement_ = 0.84, with a 95 % CI from 0.77 to 0.89; ‘number’: ICC _agreement_ = 0.78, with a 95 % CI from 0.69 to 0.85; ‘angle’: ICC _agreement_ = 0.99, with a 95 % CI from 0.98 to 0.99) and equalled 0.8 for the categorical outcome measure (‘markedness’: κ = 0.80). Again all p-values were highly significant (p < 0.001).

### 3.3 Assessment of the outcome measures

All explanatory variables selected for inclusion in the respective final models are presented in Table 3. None of the selected variables had an effect on any of the outcome measures with one exception: ‘Breed type’ had a statistically significant effect on the ‘angle’ (F_2,114_ = 8.25, p < 0.001; Fig. 2). Post-hoc tests revealed that thoroughbreds (*M* = 23.82, *SD* = 1.59) had a narrower ‘angle’ than warmbloods (*M* = 28.00, *SD* = 0.60; p = 0.040) and coldbloods (*M* = 30.98, *SD* = 0.92; p < 0.001), and that warmbloods had a narrower ‘angle’ than coldbloods (p = 0.022). Graphs for all outcome measures (‘qualitative assessment’, ‘number’, ‘angle’) grouped by explanatory variables (‘age’, ‘sex’, ‘breed type’, ‘coat colour’) can be found in Fig. 2.

**Table 3.** Statistical models, explanatory variables to be included in the final model and results for the three outcome measures.

| Outcome measure | Model | Selected explanatory variables | Test statistic | p-value |
| --- | --- | --- | --- | --- |
| Qualitative assessment | Linear mixed-effects model | sex | $F_{2,161} = 2.35$ | 0.099 |
| | | breed type | $F_{2,161} = 1.31$ | 0.273 |
| | | coat colour | $F_{3,520} = 1.12$ | 0.340 |
| Number | Linear mixed-effects model | sex | $F_{2,161} = 1.93$ | 0.149 |
| | | breed type | $F_{2,161} = 2.46$ | 0.089 |
| | | coat colour | $F_{3,516} = 0.23$ | 0.877 |
| Angle | Linear mixed-effects model | breed type | $F_{2,114} = 8.25$ | < 0.001 |

**Fig. 2.**
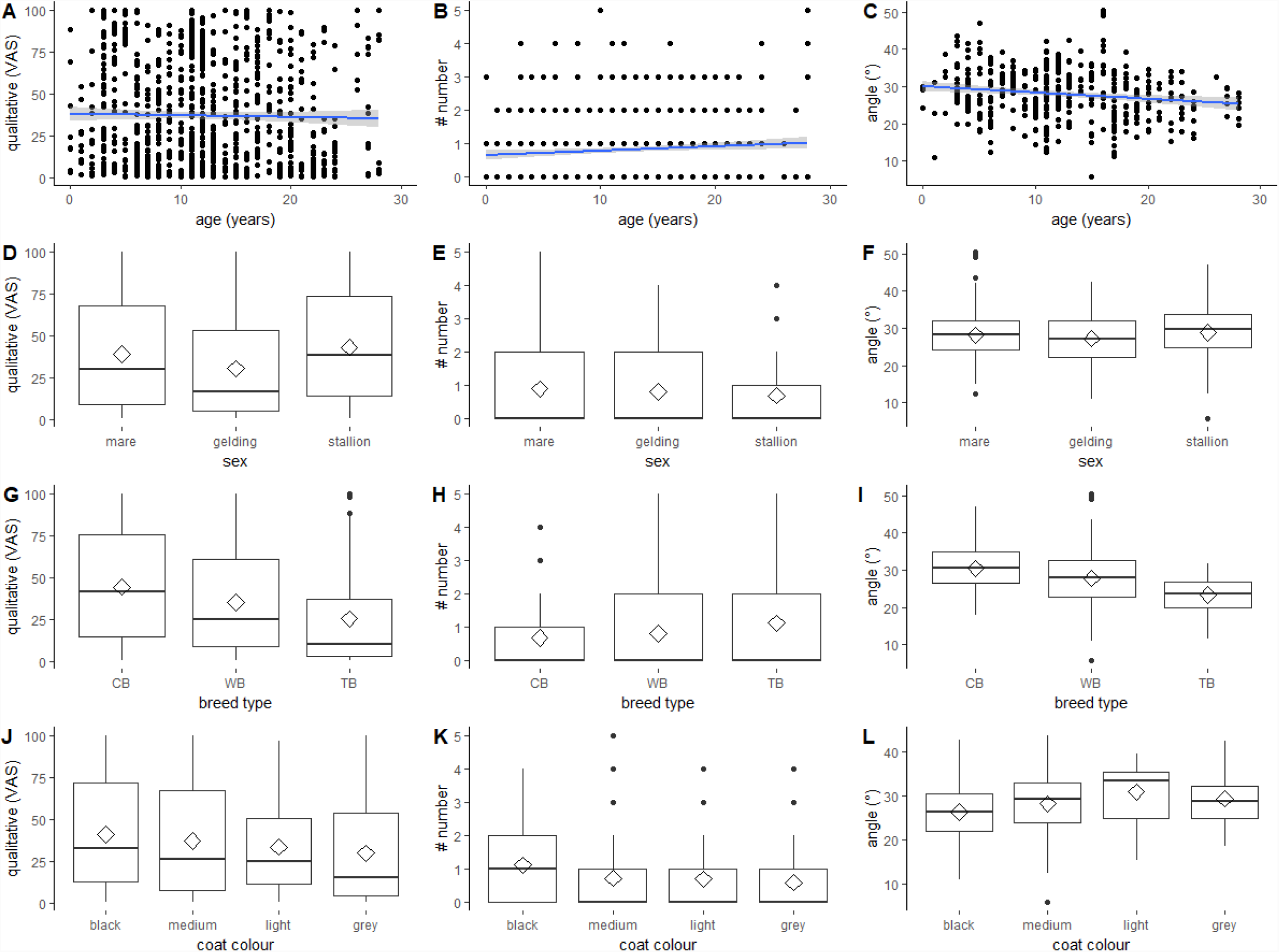
Effect of ‘age’, ‘sex’, ‘breed type’ and ‘coat colour’ on the three outcome measures. The effect of ‘age’, ‘sex’, ‘breed type’ and ‘coat colour’ on the outcome measures ‘qualitative assessment’, ‘number’ and ‘angle’. (A-C) scatter plots with regression line (method = lm, blue line) with 0.95 confidence intervals (grey). (D-L) boxplots with median (black line), mean (diamond), interquartile range (box), 1.5 x interquartile range (whiskers). (A, D, G, J) ‘qualitative assessment’ assessed on Visual Analogue Scale. (B, E, H, K) ‘number’ of wrinkles. (C, F, I, L) ‘angle’ measured in degrees. (G, H, I) ‘breed type’: coldblood (CB), warmblood (WB), thoroughbred (TB). (J, K, L) ‘coat colour’: black or dark bay (dark), medium bay (medium), light bay and palomino (light), grey and white (grey).

### 3.4 Side effects

No statistically significant effect of ‘eye’ on any of the three outcome measures was found (‘qualitative assessment’: F_1,179_ = 3.06, p = 0.082; ‘number’: F_1,353_ = 0.41, p = 0.380; ‘angle’: F_1,353_ = 3.22, p = 0.370, Fig. 3).

**Fig. 3.**
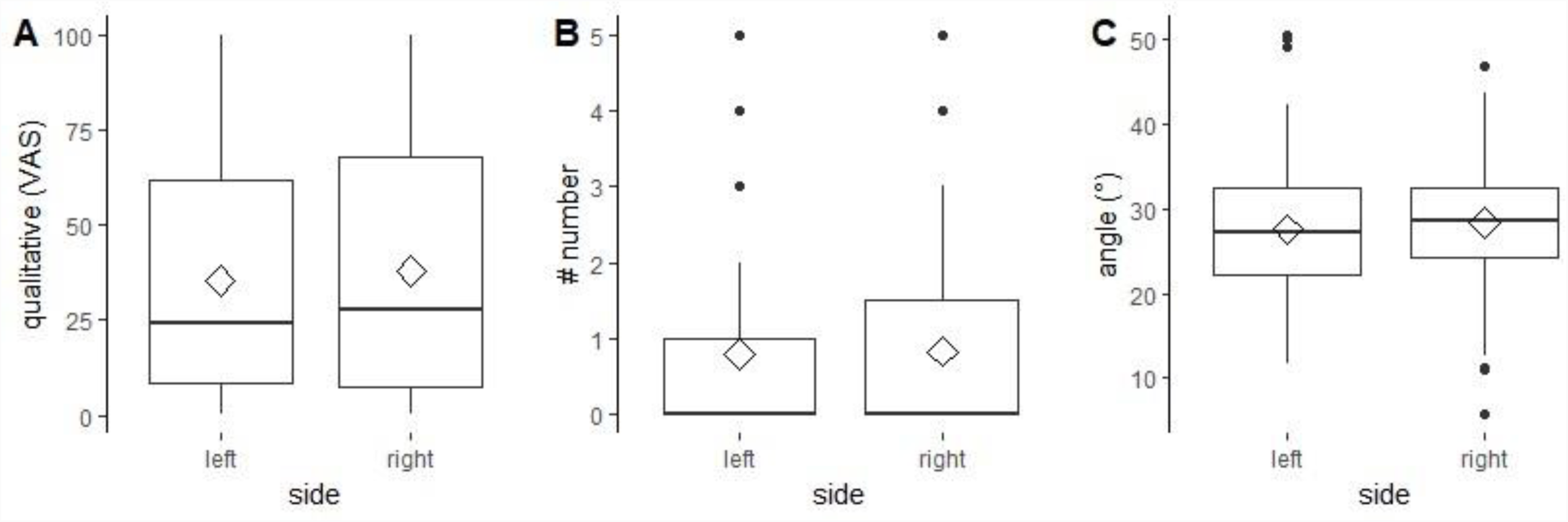
Effect of ‘side’ on the three outcome measures. The effect of ‘side’ (left, right) on the outcome measures ‘qualitative assessment’ assessed on Visual Analogue Scale (A), ‘number’ of wrinkles (B) and ‘angle’ measured in degrees (C). (A, B, C) boxplots with median (black line), mean (diamond), interquartile range (box), 1.5 x interquartile range (whiskers).

## 4 Discussion

In the present study we investigated whether ‘age’, ‘sex’, ‘breed type’, ‘Body Condition Score’ and ‘coat colour’ systematically affect eye wrinkle expression or its assessment on pictures taken from the left and the right area of horses in a presumably neutral situation. Eye wrinkle expression was assessed using five outcome measures, all of which could be assessed highly reliable with respect to both intra- and inter-observer agreement. Some outcome measures were associated, therefore only ‘qualitative assessment’, ‘number’ and ‘angle’ were further analysed. Similarly, ‘BCS’ was strongly associated with the two explanatory variables ‘sex’ and ‘breed type’ and was thus not analysed further. ‘Breed type’ influenced the width of the ‘angle’: Thoroughbreds had a narrower ‘angle’ than warmbloods and coldbloods, and warmbloods had a narrower ‘angle’ than coldbloods. The three other explanatory variables (‘age’, ‘sex’, and ‘coat colour’) did not affect any of the outcome measures, and eye wrinkle expression did not differ between the left and right eye.

### 4.1 Characteristics of the investigated sample

We found substantial variability within each outcome measure, similar to what Hintze and colleagues (2016) described for their presumably neutral control phases before the start of the experimental treatments. This variability may be explained by individual characteristics of horses independent of our tested characteristics, which did not account for the variation we found (with the exception of ‘breed type’ accounting for differences in the ‘angle’). However, other explanations need to be considered as well. First, individual horses may have reacted differently to the halter, the human handling and/or the photographing, even though we did not see indications in the horses’ behaviour supporting this assumption (in contrast to the differing reactions of the horses to the photographers and the equipment described in Hintze et al., 2016). Second, variation in eye wrinkle expression across horses could be caused by differences in underlying mood states, but this explanation is speculative since we did not assess mood in the present study.

Certain confounding effects (e.g. between ‘breed type’ and ‘farm’, see below) could not be ruled out since our sample was not fully balanced across all explanatory variables. However, we counteracted this limitation as much as possible under non-experimental conditions by ensuring a relatively large sample for the different categories of ‘sex’ (70 mares, 62 geldings, and 49 stallions), ‘breed type’ (52 coldbloods, 104 warmbloods, and 17 thoroughbreds) and ‘age’ (including young to very old horses).

### 4.2 Adaptation of the eye wrinkle assessment scale with respect to intra- and inter-observer agreement

Our eye wrinkle assessment scale is based on the scale described by Hintze et al. (2016) but was adapted with respect to both the outcome measures being included (‘eyebrow raised’) or excluded (‘eyelid shape’, ‘eye white’) as well as the definition and measurement of one outcome measure (‘qualitative assessment’). ‘Eye white’ tended to be systematically affected by the emotional situations induced in the study by Hintze et al. (2016) with more horses showing ‘eye white’ in the negative and fewer horses showing ‘eye white’ in the positive situations compared to control phases. Therefore, we recommend to include ‘eye white’ (either as a binary measure or as ‘percentage of visible white of the total visible eye area’ as it has been assessed in dairy cows, e.g. Sandem et al., 2002) as an outcome measure in future studies investigating the effect of different emotional situations on changes in the eye area.

Results for intra- and inter-observer agreement in our study were largely comparable with the results from Hintze and colleagues, with ‘excellent’ (continuous measures, ICC > 0,75) or ‘almost perfect’ agreement (categorical measure, κ > 0.8) for all outcome measures in the present study. Whereas inter-observer agreement for ‘eyelid shape’ was only moderate in the study by Hintze et al. (2016), agreement for our replacing outcome measure ‘eyebrow raised’ was ‘excellent’ (ICC > 0,75). ‘Eyebrow raised’ was assessed as a continuous measure on a Visual Analogue Scale and not on an ordinal scale with three distinct categories, which had previously been used for ‘eyelid shape’. Besides potentially being more sensitive, VAS have shown to improve inter-observer agreement and to be preferred by raters compared to a 3-point ordinal scale (Tuyttens et al., 2009). We cannot disentangle whether our improved inter-observer agreement was due to the adapted measure *per se* or the changed scoring method, but both raters preferred the VAS to the ordinal scales when using it for ‘eyebrow raised’ and ‘qualitative assessment’, a preference that is in line with what Tuyttens and colleagues report (2009).

### 4.3 Relationship between ‘Body Condition Score’ and the two explanatory variables ‘breed type’ and ‘sex’

Variation in ‘BCS’ could be explained by the three ‘breed types’, with coldbloods having the highest scores, followed by warmbloods and thoroughbreds. This finding is consistent with what has been reported by Giles et al. (2014), who found that breed was the risk factor most strongly associated with obesity in horses. In line with this, Visser et al. (2014) found that coldbloods were more prone to develop a higher body condition score compared to thoroughbreds, which was also the case in our study. However, the association between ‘BCS’ and ‘breed type’ in our study could also be explained by confounds in our sample since most coldbloods were from Farm 7 and the management practices on a farm, especially the feeding regime, including the amount of feed and its nutritional value, can influence the body condition of horses. This confound may also explain the strong association between ‘sex’ and ‘breed type’ since most of the stallions on Farm 7 were coldbloods.

### 4.4 Interpretation of the results of the outcome measures

‘Breed type’ systemically affected the ‘angle’ between the extensions of a line through the eyeball and a line through the topmost wrinkle. ‘Angle’ is a measure of muscle contraction of the inner eyebrow raiser with a wider angle reflecting a stronger contraction and a narrower angle reflecting a comparably more relaxed muscle. In our study, thoroughbreds had the narrowest ‘angle’, followed by warmbloods and coldbloods. This result may simply reflect differences in the degree of contraction of the underlying muscle between ‘breed types’ but other explanations need to be discussed as well. Confounders might have affected our result with most coldbloods being derived from Farm 7. However, thoroughbreds came from six and warmbloods from all seven farms, which is why the confounding of ‘breed type’ and ‘farm’ cannot fully explain the variation in ‘angles’. Another confound that needs to be considered is the one between ‘breed type’ and the different housing conditions of the horses. Whereas most thoroughbreds were kept either in groups, paddock boxes or boxes with daytime turnout, most of the coldbloods were housed in standard single horse boxes. Housing conditions might have affected mood and thus eye wrinkle expression with group-housed horses and horses with more space as well as physical contact to conspecifics having a more relaxed inner eyebrow raiser than horses kept in single boxes. However, our study was neither designed to study the effect of housing conditions on eye wrinkle expression, nor was mood investigated. Thus, these explanations remain speculative and further research is needed to study the relationship between housing conditions, long-lasting mood states and eye wrinkle expression.

Beside the systematic effect of ‘breed type’ on ‘angle’, there was no further effect of any of the explanatory variables on our outcome measures. This finding indicates that eye wrinkle expression can be assessed regardless of ‘age’, ‘sex’ and ‘coat colour’, while ‘breed type’ should be considered in future studies. Our study does not give further insight into the relationship between emotion or mood and eye wrinkle expression, but it shows that eye wrinkle expression in horses cannot simply be explained by horses’ characteristics such as ‘age’, ‘sex’, ‘coat colour’ or ‘breed type’ (with the exception of the ‘angle’ for the latter).

### 4.5 Side effects

In our study no difference in eye wrinkle expression between the left and right eye area was found in a neutral situation, neither did Hintze and colleagues find a difference in positively and negatively valenced situations. We surmise that eye wrinkle expression in horses is not affected by emotionally induced or morphological asymmetry in the face, determined by, for example, facial soft tissue (Sackheim, 1985). The results from both studies suggest that it is irrelevant which eye area is assessed; this could be advantageous if eye wrinkle expression is used in the future as an on-farm indicator of horses’ emotional states.

## 5 Conclusion

To our knowledge, this was the first study systematically investigating the effect of individual characteristics on eye wrinkle expression and its assessment in horses. We conclude that our eye wrinkle assessment scale can be used reliably and regardless of horses’ age, sex, coat colour and breed type (here with the exception of the ‘angle’). Thus, the adapted scale is a promising tool to assess eye wrinkles in horses, but to what extent these are systematically affected by mood or emotion needs further investigation and validation.

## Author Contributions

LS, KK and SH conceived the study. LS and SH developed the methodology. LS (and SH) collected the data. LS scored all pictures and performed the data analysis. LS and SH wrote the manuscript. All authors edited the manuscript, contributed to manuscript revision, read and approved the submitted version.

## Acknowledgments

We would like to thank the horse handlers Alisa Herbst and Valeria Gift for their enthusiasm and patience in working with us, and Samantha Smith for picture taking and processing on Farm 7. We are grateful to all the horse owners who kindly allowed us to photograph their horses for this study. Moreover, we would like to thank Christoph Winckler for his support and comments on this manuscript, as well as for valuable discussions.

## References

Akazaki, S., Nakagawa, H., Kazama, H., Osanai, O., Kawai, M., Takema, Y., Imokawa, G., 2002. Age-related changes in skin wrinkles assessed by a novel three-dimensional morphometric analysis. Br. J. Dermatol. 147, 689–695.

André, C.D.S., Narula, S.C., Elian, S.N., Tavares, R.A., 2003. An overview of the variables selection methods for the minimum sum of absolute errors regression. Stat. Med. 22, 2101–2111.

Broadhurst, D., Goodacre, R., Jones, A., Rowland, J.J., Douglas, B.K., 1997. Genetic algorithms as a method for variable selection in multiple linear regression and partial least squares regression, with applications to pyrolsis mass spectrometry. Anal. Chim. Acta.

Caeiro, C.C., Burrows, A.M., Waller, B.M., 2017. Development and application of CatFACS: Are human cat adopters influenced by cat facial expressions? Appl. Anim. Behav. Sci. 189, 66–78.

Caeiro, C.C., Waller, B.M., Zimmermann, E., Burrows, A.M., Davila-Ross, M., 2013. OrangFACS: A muscle-based facial movement coding system for orangutans (Pongo spp.). Int. J. Primatol. 34, 115–129.

Cicchetti, D. V, 1994. Guidlines, criteria, and rules of thumb for evalauting normed and standardized assessment instruments in psychology. Psychol. Assess. 6, 284–290.

Corel Cooperation, 2014. CorelDRAW Version 17.1.0.572.

Cramér, H., 2016. Mathematical methods of statistics (PMS-9). Princeton university press.

Dalla Costa, E., Minero, M., Lebelt, D., Stucke, D., Canali, E., Leach, M.C., 2014. Development of the Horse Grimace Scale (HGS) as a pain assessment tool in horses undergoing routine castration. PLoS One 9, 1–10.

Di Giminiani, P., Brierley, V.L.M.H., Scollo, A., Gottardo, F., Malcolm, E.M., Edwards, S.A., Leach, M.C., 2016. The assessment of facial expressions in piglets undergoing tail docking and castration: Toward the development of the piglet grimace scale. Front. Vet. Sci. 3, 1–10.

Ekman, P., Friesen, W. V, 1976. Measuring facial movement. Environ. Psychol. Nonverbal Behav. 1, 56–75.

Gamer, M., Lemon, J., Fellows, I., Singh, P., 2012. irr: Various coefficients of interrater reliability and agreement. R package version 0.84. https://CRAN.R-project.org/package=irr.

Giles, S.L., Rands, S.A., Nicol, C.J., Harris, P.A., 2014. Obesity prevalence and associated risk factors in outdoor living domestic horses and ponies. PeerJ 2, e299.

Gleerup, K.B., Forkman, B., Lindegaard, C., Andersen, P.H., 2015. An equine pain face. Vet. Anaesth. Analg. 42, 103–114.

Guesgen, M.J., Beausoleil, N.J., Leach, M., Minot, E.O., Stewart, M., Stafford, K.J., 2016. Coding and quantification of a facial expression for pain in lambs. Behav. Processes 132, 49–56.

Henneke, D.R., Potter, G.D., Kreider, J.L., Yeates, B.F., 1983. Relationship between condition score, physical measurements and body fat percentage in mares. Equine Vet. J. 15, 371–372.

Hintze, S., Smith, S., Patt, A., Bachmann, I., Würbel, H., 2016. Are eyes a mirror of the soul? What eye wrinkles reveal about a horse’s emotional state. PLoS One 11, 1–15.

Keating, S.C.J., Thomas, A.A., Flecknell, P.A., Leach, M.C., 2012. Evaluation of EMLA cream for preventing pain during tattooing of rabbits: Changes in physiological, behavioural and facial expression responses. PLoS One 7, 1–11.

Koo, T.K., Li, M.Y., 2016. A guideline of selecting and reporting intraclass correlation coefficients for reliability research. J. Chiropr. Med. 15, 155–163.

Kuhn, M., Contributions from Wing, J., Weston, S., Williams, A., Keefer, C., Engelhardt, A., Cooper, T., Mayer, Z., Kenkel, B., the R Core Team, T., Benesty, M., Lescarbeau, R., Ziem, A., Scrucca, L., Tang, Y., Candan, C., Hunt, T., 2018. caret: Classification and regression training. R package version 6.0-80. https://CRAN.R-project.org/package=caret.

Kwon, Y.H., da Vitoria Lobo, N., 1999. Age classification from facial images. Zhurnal Eksp. i Teor. Fiz. 74, 1–21.

Landis, J.R., Koch, G.G., 1977. The measurement of observer agreement for categorical data. Biometrics 159–174.

Langford, D.J., Tuttle, A.H., Brown, K., Deschenes, S., Fischer, D.B., Mutso, A., Root, K.C., Sotocinal, S.G., Stern, M.A., Mogil, J.S., 2010. Social approach to pain in laboratory mice. Soc. Neurosci. 5, 163–170.

Lansade, L., Foury, A., Reigner, F., Vidament, M., Guettier, E., Bouvet, G., Soulet, D., Parias, C., Ruet, A., Mach, N., 2018. Progressive habituation to separation alleviates the negative effects of weaning in the mother and foal. Psychoneuroendocrinology 97, 59–68.

Lumley, T., Alan, M., 2009. leaps: regression subset selection. R package version 2.9. https://CRAN.R-project.org/package=leaps.

Marsh-Richard, D.M., Hatzis, E.S., Mathis, C.W., Venditti, N., Dougherty, D.M., 2009. Adaptive Visual Analog Scales (AVAS): A modifiable software program for the creation, administration, and scoring of visual analog scales. Behav. Res. Methods 41, 99–106.

Martin, P., Bateson, P.P.G., 2007. Measuring behaviour: an introductory guide. Cambridge University Press, Cambridge.

McLennan, K.M., Rebelo, C.J.B., Corke, M.J., Holmes, M.A., Leach, M.C., Constantino-Casas, F., 2016. Development of a facial expression scale using footrot and mastitis as models of pain in sheep. 1–32.

Parr, L., Waller, B., 2006. The evolution of human emotion. In: Evolution of Nervous Systems: A Comprehensive Reference. Academic Press Inc.

Parr, L.A., Waller, B.M., Burrows, A.M., Gothard, K.M., Vick, S.J., 2010. Brief communication: MaqFACS: A muscle-based facial movement coding system for the rhesus macaque. Am. J. Phys. Anthropol. 143, 625–630.

Parr, L.A., Waller, B.M., Vick, S.J., 2007a. New developments in understanding emotional facial signals in chimpanzees. Curr. Dir. Psychol. Sci. 16, 117–122.

Parr, L.A., Waller, B.M., Vick, S.J., Bard, K.A., 2007b. Classifying chimpanzee facial expressions using muscle action. Emotion 7, 172–181.

Patton, F.J., Campbell, P.E., 2011. Using eye and profile wrinkles to identify individual white rhinos. Pachyderm - J. African Elephant, African Rhino Asian Rhino Spec. Groups 84–86.

Paul, E.S., Harding, E.J., Mendl, M., 2005. Measuring emotional processes in animals: The utility of a cognitive approach. Neurosci. Biobehav. Rev. 29, 469–491.

Pinheiro, J., Bates, D., DebRoy, S., Sarkar, D., Core Team, R T., 2018. nlme: Linear and Nonlinear Mixed Effects Models_. R package version 3.1-137, URL: https://CRAN.R-project.org/package=nlme.

R Core Team, 2014. R: A language and environment for statistical computing. R Foundation for Statistical Computing, Vienna, Austria. URL http://www.R-project.org/.

RStudio Team, 2016. RStudio: Integrated development for R. RStudio, Inc., Boston, MA URL http://www.rstudio.com/.

Sandem, A.I., Braastad, B.O., Bøe, K.E., 2002. Eye white may indicate emotional state on a frustration-contentedness axis in dairy cows. Appl. Anim. Behav. Sci. 79, 1–10.

Sotocinal, S.G., Sorge, R.E., Zaloum, A., Tuttle, A.H., Martin, L.J., Wieskopf, J.S., Mapplebeck, J.C.S., Wei, P., Zhan, S., Zhang, S., McDougall, J.J., King, O.D., 2011. The Rat Grimace Scale: A partially automated method for quantifying pain in the laboratory rat via facial expressions. Mol. Pain 7, 55.

Tu, Y.K., Kellett, M., Clerehugh, V., Gilthorpe, M.S., 2005. Problems of correlations between explanatory variables in multiple regression analyses in the dental literature. Br. Dent. J. 199, 457–461.

Tuyttens, F.A.M., Sprenger, M., Van Nuffel, A., Maertens, W., Van Dongen, S., 2009. Reliability of categorical versus continuous scoring of welfare indicators: Lameness in cows as a case study. In: Animal Welfare.

v. Lenth, R., 2016. Least-squares means: The R package lsmeans. J. Stat. Softw. 69, 1–33.

Venables, W.N., Ripley, B.D., 2002. Modern Applied Statistics with S, 4th ed. Springer, New York.

Vick, S.J., Waller, B.M., Parr, L.A., Pasqualini, M.C.S., Bard, K.A., 2007. A cross-species comparison of facial morphology and movement in humans and chimpanzees using the Facial Action Coding System (FACS). J. Nonverbal Behav. 31, 1–20.

Visser, E.K., Neijenhuis, F., de Graaf-Roelfsema, E., Wesselink, H.G.M., de Boer, J., van Wijhe-Kiezebrink, M.C., Engel, B., van Reenen, C.G., 2014. Risk factors associated with health disorders in sport and leisure horses in the Netherlands. J. Anim. Sci. 92, 844–855.

Waller, B., Caeiro, C.C., Peirce, K., Burrows, A., Kaminski, J., 2013. DogFACS: the dog facial action coding system.

Waller, B.M., Lembeck, M., Kuchenbuch, P., Burrows, A.M., Liebal, K., 2012. GibbonFACS: A muscle-based facial movement coding system for hylobatids. Int. J. Primatol. 33, 809–821.

Wathan, J., Burrows, A.M., Waller, B.M., McComb, K., 2015. EquiFACS: The equine facial action coding system. PLoS One 10, 1–35.

Wemelsfelder, F., 1997. The scientific validity of subjective concepts in models of animal welfare. Anim. Stud. Repos. 53, 75–88.

Wewers, M.E., Lowe, N.K., 1990. A critical review of visual analogue scales in the measurement of clinical phenomena. Res. Nurs. Health 13, 227–236.

